# Assembly-based computations through contextual dendritic gating of plasticity

**DOI:** 10.1101/2025.07.22.666089

**Authors:** Sebastian Onasch, Christoph Miehl, M. Maurycy Miękus, Julijana Gjorgjieva

## Abstract

Neuronal assemblies – groups of strongly connected neurons – are considered the basic building blocks of perception and memory in the brain by encoding representations of specific concepts. Despite recent evidence for the biological basis behind the existence and formation of such assemblies, computational models often fall short of showing how assemblies can be flexibly learned and combined to perform real-world computations. A prominent problem is ‘catastrophic forgetting’, where learning a new assembly can disrupt existing connectivity structure and lead to forgetting previously learned assemblies. We propose a biologically plausible computational model, where dendritic compartments (instead of neurons) are the loci for learning and inhibition gates learning in a dendrite-specific manner, to flexibly learn new stimuli without forgetting of old ones. By learning stable projections from one brain region into another and associations between different brain regions, we demonstrate how the proposed assembly framework implements the basic building blocks for diverse computations. In a visual-auditory association task, we demonstrate how the context-specific assembly computations can be used to correctly separate ambiguous stimuli based on their dendritic representations. Our models provide unique insights and predictions for how hierarchically connected brain areas use their biological components to implement flexible yet robust learning.

## Introduction

The brain can represent, flexibly learn, and stably store information. A long-standing hypothesis is that groups of neurons with highly correlated activity, known as neuronal ensembles, provide the basis for representations in the brain (Buzsáki, 2010; Yuste, 2015; Eichenbaum, 2018). Groups of strongly connected neurons called neuronal assemblies are hypothesized to be the structural basis of these correlated ensembles (Miehl et al., 2023). A core assumption is that each assembly represents a specific concept or feature, acting as a fundamental unit for storing memories (Neves et al., 2008). Even though many experimental studies have supported the idea of ensembles as the basic memory unit (Yuste, 2015; Josselyn and Tonegawa, 2020), past theoretical studies were either based on over-simplified network structures (Hopfield, 1982) or have mainly focused on how assembly structures can be learned via synaptic plasticity mechanisms (Clopath et al., 2010; Litwin-Kumar and Doiron, 2014; Zenke et al., 2015; Wu et al., 2020; Montangie et al., 2020). Only a few studies have shown that assemblies can be flexibly learned and combined to perform interesting computations (Tetzlaff et al., 2015; Papadimitriou et al., 2020; Weidel et al., 2021). Despite this interest and prior work on assemblies, several challenges remain: 1) overlaps in assemblies in terms of participating neurons often lead to the merging of assembly structures during ongoing plasticity, and 2) imprinting a new assembly into a network with an already existing connectivity structure can lead to ‘forgetting’ of stored representations.

At the same time, artificial neural networks (ANNs) have proved very successful and often exceed humans in solving specific tasks (Silver et al., 2017). However, ANNs mainly rely on assumptions that are not always biologically plausible, and learning new or additional tasks in ANNs leads to decreased performance and catastrophic forgetting in the worst case (Parisi et al., 2019). While multiple solutions have been suggested to counteract forgetting (Kirkpatrick et al., 2017; Masse et al., 2018; Jedlicka et al., 2022; Mei et al., 2022; Zenke and Laborieux, 2024), here we show how this problem can be solved in hierarchical recurrent neural networks using two specific biological components: nonlinear dendritic compartments and inhibitory context-dependent gating. We propose that dendritic compartments (instead of neurons) are the loci for learning, and inhibition gates learning in a dendrite-specific manner.

First, we introduce dendritic compartments with nonlinear integration properties to each neuron (Fig. 1A). Recent computational studies have used dendrites for feature binding (Legenstein and Maass, 2011), multi-task learning (Iyer et al., 2022; Wybo et al., 2023), linking of memories (Kastellakis et al., 2016, 2023), solving the stability-plasticity problem (Wilmes and Clopath, 2023), or exploring the interplay of somatic and dendritic inhibition (Pedrosa and Clopath, 2020) (as reviewed in Payeur et al. (2019); Poirazi and Papoutsi (2020)). Furthermore, dendritic compartments have been successfully implemented in machine learning applications to solve various tasks such as the credit assignment problem in deep neuronal networks (Guerguiev et al., 2017; Sacramento et al., 2018; Payeur et al., 2021; Greedy et al., 2022; Galloni et al., 2025), catastrophic forgetting (Masse et al., 2018; Grewal et al., 2021; Iyer et al., 2022; Wybo et al., 2023), parameter-efficient learning (Chavlis and Poirazi, 2025), have been proposed to solve classic machine learning tasks (Jones and Kording, 2020), and more efficient neuromorphic computing (Bhaduri et al., 2018) (as reviewed in Acharya et al. (2022); Pagkalos et al. (2024)).

**Figure 1.**
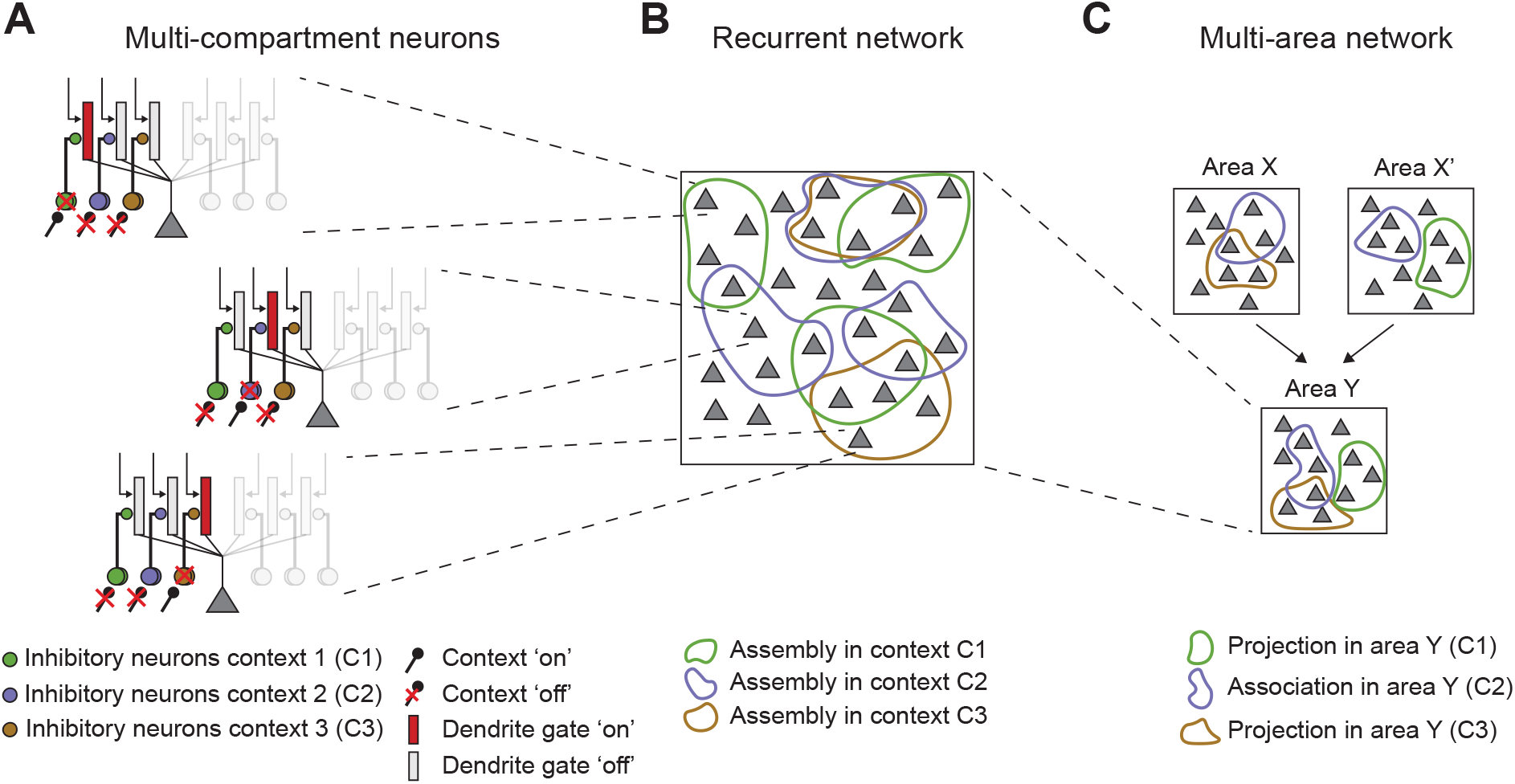
Bridging spatial scales from local dendritic properties to multi-area computations. **A**. Schematic of multicompartment neurons with contextual dendrite-specific inhibitory control. A dendrite is gated ‘on’ if the inhibitory neurons specific to a given context are inhibited, and a dendrite is gated ‘off’ if the inhibitory neurons are not inhibited. **B**. Overlapping assembly structures can be learned in a recurrent network without forgetting previously learned ones, where a neuron can only participate in a given assembly with a dendrite that is gated ‘on’ in the appropriate context. **C**. Assembly computations in a multiarea network.

Second, we include inhibition as a mechanism for dendrite-specific gating. Cortical circuits contain a diversity of inhibitory cell types known to play specific roles in regulating cortical function and stability (Pfeffer et al., 2013). For example, parvalbumin (PV) expressing inhibitory neurons stabilize excitatory dynamics (Veit et al., 2017; Sanzeni et al., 2020; Lagzi et al., 2021; Festa et al., 2024; Palmigiano et al., 2023) while somatostatin (SOM) expressing inhibitory neurons modulate the circuit (Adesnik et al., 2012; Bos et al., 2025). Motivated by the top-down control of vasoactive intestinal peptide (VIP) expressing inhibitory neurons and the disinhibitory VIP-SOM circuit (Pi et al., 2013; Canto-Bustos et al., 2022; Waitzmann et al., 2024), we incorporate inhibition as a signal to gate dendrites ‘on’ or ‘off’. We interpret (dis-) inhibition as a ‘context’ signal, supporting dendrite-specific control of synaptic plasticity (Fig. 1A). Context-dependent gating of activity has been utilized in ANNs for multitask learning (Wybo et al., 2023), extending the dynamical regime of ANNs (Krishnamurthy et al., 2022), and supporting high-capacity memory networks (Podlaski et al., 2025).

We show that adding two biological constraints, dendritic nonlinear integration and dendritic gating via inhibition, to hierarchically organized recurrent network models enables the learning of stable overlapping assemblies without forgetting previously learned features (Fig. 1B). We demonstrate the potential of this framework to perform assembly computations based on projections and associations across multiple areas while amplifying information through recurrent connectivity and separating ambiguous stimuli (Fig. 1C). Therefore, by providing a biologically grounded solution to the problem of catastrophic forgetting, we open the door to the mechanistic implementations of other forms of associative memory formation, and provide a basis for flexible yet robust learning in ANNs.

## Results

### Inhibitory context gates dendrite-specific excitatory long-term plasticity

To investigate the learning of stable but flexible assemblies that encode distinct concepts in recurrent neural networks with spiking neurons, we identified two biologically motivated components as key: dendritic nonlinearities and context-specific inhibitory gating. We modeled each excitatory neuron as a multi-compartment unit, with a somatic leaky-integrate-and-fire compartment connected to multiple dendritic branches (Fig. 2A). The neurons receive excitatory inputs to their dendrites via excitatory synapses with two types of ion channels, AMPA and NMDA, which integrate inputs linearly and nonlinearly, respectively (Methods). Therefore, strong excitatory inputs can induce plateau potentials (‘NMDA spikes’), which last several milliseconds and have been observed experimentally (Schiller et al., 2000). We also modeled inhibitory synapses so that sufficiently strong inhibition can prevent these NMDA spikes, in line with experimental studies showing that inhibition can operate on single dendritic branches (Gidon and Segev, 2012; Stokes et al., 2014) and can control NMDA receptor activation (Schulz et al., 2018; Davenport et al., 2021).

**Figure 2.**
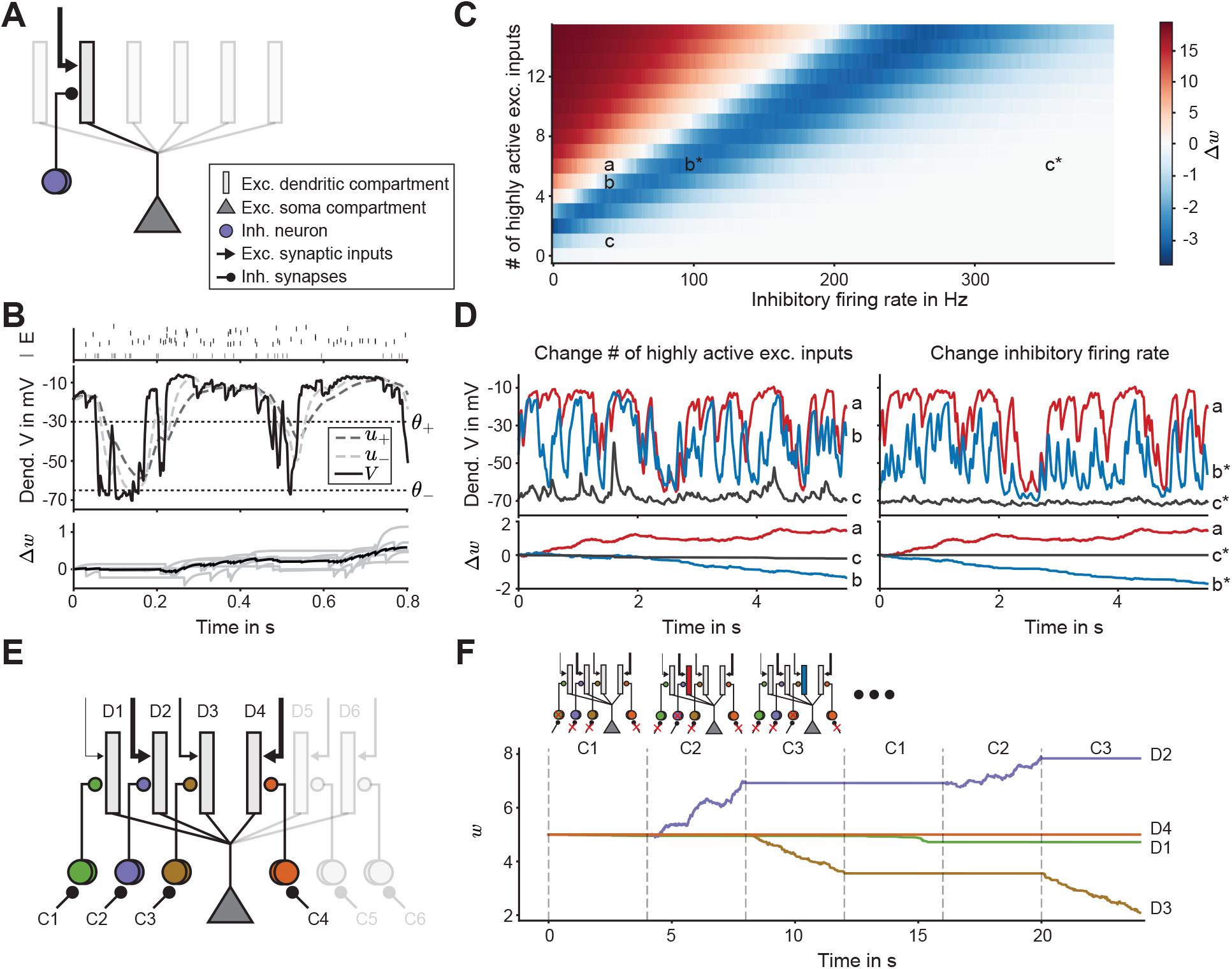
Gating of excitatory plasticity via inhibitory context signals at nonlinear dendrites. **A**. Schematic of a single excitatory neuron with a somatic compartment and six dendritic compartments. Each dendrite receives excitatory and inhibitory input (only shown for one dendrite). **B**. Inhibitory and excitatory presynaptic spikes (top). When the low-pass filtered dendritic voltage (*u*_−_, middle) at a synapse exceeds the LTD threshold (*θ*_−_, middle) at the time of a presynaptic spike, the synaptic weight (bottom) undergoes LTD. If the low-pass filtered dendritic voltage (*u*_+_, middle) with a longer time constant exceeds the threshold *θ*_−_ at the same time as the momentary dendritic voltage (*V*) surpasses a higher threshold (*θ*_+_), the synaptic weight undergoes LTP at all times (Methods). Mean (black) vs. individual (gray) synaptic weight change of highly active excitatory input synapses (bottom). **C**. Mean synaptic weight change Δ*w* as a function of the number of highly active excitatory synapses and the firing rate of inhibitory neurons. Long-term depression (LTD) is indicated by shades of blue, and long-term potentiation (LTP) by shades of red. **D**. Left: Filtered dendritic voltage *u*_+_ (top), and mean synaptic weight change Δ*w* (bottom) for a different number of highly active excitatory inputs (6 – red, 5 – blue, 1 – gray). Right: Same as left, for changing the inhibitory firing rate (40 Hz – red, 100 Hz – blue, 350 Hz – gray). Letters (a, b, c, a*, b*, c*) correspond to cases in panel C. **E**. Schematic of a single excitatory neuron with six dendritic compartments (D1-D6). Each dendrite receives a different number of highly active excitatory input synapses (indicated by the thickness of arrows) and a separate inhibitory input. Each inhibitory population is controlled by a separate inhibitory context signal (C1-C6). See also Fig. 1A. **F**. Mean weight (*w*) change at dendrite D1, D2, D3, and D4. The activity of the respective inhibitory group is reduced by a context signal (C1, C2, and C3) for 4 seconds (gray dashed lines). D1 receives 1, D2 receives 6, D3 receives 4 and D4 receives 7 highly active excitatory inputs. The inhibitory firing rate is 400 Hz if a context is ‘off’ and 0 Hz if a context is ‘on.’ Dendrites color-coded in red undergo LTP, in blue LTD, and in gray no plasticity.

To form stimulus-specific assemblies, we implemented long-term synaptic plasticity at excitatory synapses, which modifies excitatory synapses in an activity-dependent manner. Synaptic change follows a dendritic voltage-dependent plasticity rule where the sign of induced plasticity (long-term potentiation, LTP, vs. depression, LTD) depends on the low-pass filtered dendritic voltage traces relative to a plasticity threshold (Clopath et al., 2010; Bono and Clopath, 2017) (Fig. 2B, Methods). A synapse undergoes LTD if the dendritic voltage is slightly depolarized at the time of the presynaptic spike (above a threshold *θ*_−_), whereas it undergoes LTP if depolarized strongly (above a second threshold *θ*_+_). Under these conditions, the balance of excitatory and inhibitory inputs determines the sign and amount of synaptic plasticity at the dendrite (Fig. 2C). For example, in the model, many active excitatory inputs can repeatedly evoke NMDA spikes when inhibition is weak, and therefore, induce LTP (Fig. 2C, D). Decreasing the number of highly active excitatory inputs can shift the plasticity regime to LTD, and even to no plasticity if only few inputs are active (Fig. 2D, left). Equivalently, increasing the inhibitory firing rate can also shift the plasticity regime to LTD and eventually to no plasticity (Fig. 2D, right). An important factor enabling the dynamic switch between different regimes, LTP, LTD and no plasticity, is the nonlinearity of the NMDA synapses. With linear NMDA synapses, switches between plasticity regimes require much more drastic changes of excitatory and inhibitory input parameters (Suppl. Fig. S1). Each dendritic branch in our model can be considered an independent component, separate from the activity and plasticity dynamics at the other dendrites of the same neuron. Experimental studies have found such branch-specificity in the visual cortex (Jia et al., 2010), motor cortex (Cichon and Gan, 2015; Kerlin et al., 2019), and the hippocampus (Rashid et al., 2020; Moore et al., 2022).

Inhibitory neurons have been shown to modulate dendritic excitability (Bilash et al., 2023) and to gate synaptic plasticity (Canto-Bustos et al., 2022; Pi et al., 2013; Adesnik et al., 2012). Inspired by these experimental findings, we included inhibitory neurons that each represent a specific context (C1-C6), which can be switched ‘on’ one at a time (Fig. 2E). We assumed that each dendritic compartment of the excitatory neurons receives inhibitory input from a distinct context. When the context is ‘off’, the inhibitory input is active, and the inputs targeting this dendrite experience no plasticity. When a given context is ‘on’, the corresponding inhibitory input is inactive, the dendrite is disinhibited and can be shifted into different plasticity regimes (LTP, LTD or no plasticity) depending on the number of highly active excitatory inputs it receives. Hence, the baseline state is one with highly active inhibitory neurons preventing ongoing plasticity. For example, when context C1 is ‘on’, there is barely any plasticity on the corresponding dendrite D1 if it receives few highly active inputs (Fig. 2F). If context C2 is ‘on’ and the corresponding dendrite D2 receives many highly active excitatory inputs, then the connections to D2 undergo LTP. Finally, if context C3 is on, but the corresponding dendrite D3 receives an intermediate number of highly active inputs, the connections to D3 experience LTD. Importantly, even if a dendrite receives many strong excitatory inputs, it does not undergo any plasticity if the context signal remains ‘off’ (D4). Hence, while dendritic depolarization determines the direction of plasticity in our model, somatic spiking only minimally affects plasticity in line with experimental findings (Gambino et al., 2014; Golding et al., 2002; Cichon and Gan, 2015).

Hence, we have proposed a biologically grounded modeling framework to capture how excitatory plasticity induced by nonlinear NMDA spikes can be flexibly gated ‘on’ or ‘off’ in a dendrite-specific manner by inhibitory context signals. We demonstrated that if a dendrite is gated ‘on,’ the type of plasticity experienced by the synapses on the dendrite (no plasticity, LTD, or LTP) depends on the number of highly active excitatory inputs that arrive at the dendrite, providing a well controlled substrate for learning.

### Nonlinear dendrites and inhibitory context lead to the formation of dendrite-specific assemblies

We next combined the two biologically motivated components, nonlinear dendritic integration and inhibition-specific gating, with biologically inspired synaptic plasticity to stably learn assemblies that encode percepts or stimulus features in a recurrently connected network of excitatory neurons. To investigate the mutual potentiation of connections between neurons that underlie the assemblies, we connected 400 multi-compartment excitatory neurons in an all-to-all manner with initially weak but plastic connections on the dendrites (Fig. 3A, Methods). To stimulate the network, we drove each neuron with input from a random set of excitatory feedforward neurons on its dendrites. Thus, active feedforward inputs strongly activate a subset of neurons in the recurrent network (Fig. 3B, top and middle). We implemented activity-dependent plasticity at the synaptic connections from feed-forward and recurrent inputs at the dendrites. The type of plasticity at each synapse (LTP, LTD or no plasticity) depends on the number of highly active feedforward and recurrent excitatory inputs received at each dendrite and whether the inhibitory context for that dendrite is ‘on’ (Fig. 2). Assuming the same context (C1) is always ‘on’, we observed the emergence of an assembly of strongly mutually connected neurons in the recurrent network, as seen in the neuron-neuron connectivity matrix (Fig. 3C, left). Neurons with ‘on’-gated dendrites, which receive many active inputs, acquire the strongest mutual connections, as seen in the dendrite-neuron connectivity matrix (Fig. 3C, right). With this plasticity mechanism, only the weights that are part of the assembly (‘within’) increase. In contrast, weights from other neurons to the assembly neurons (‘into’), weights from assembly neurons to other neurons (‘from’) and weights between other neurons (‘outside’) do not change or decrease, as they receive fewer excitatory inputs or too much dendritic inhibition (Fig. 3B, bottom). In addition to the inhibitory context, the network receives feedforward and recurrent inhibition that, together with a biologically motivated dendrite-specific normalization mechanism, support the formation of refined assemblies while controlling assembly size (Suppl. Fig. S2, see Supplementary Text 1 for details).

**Figure 3.**
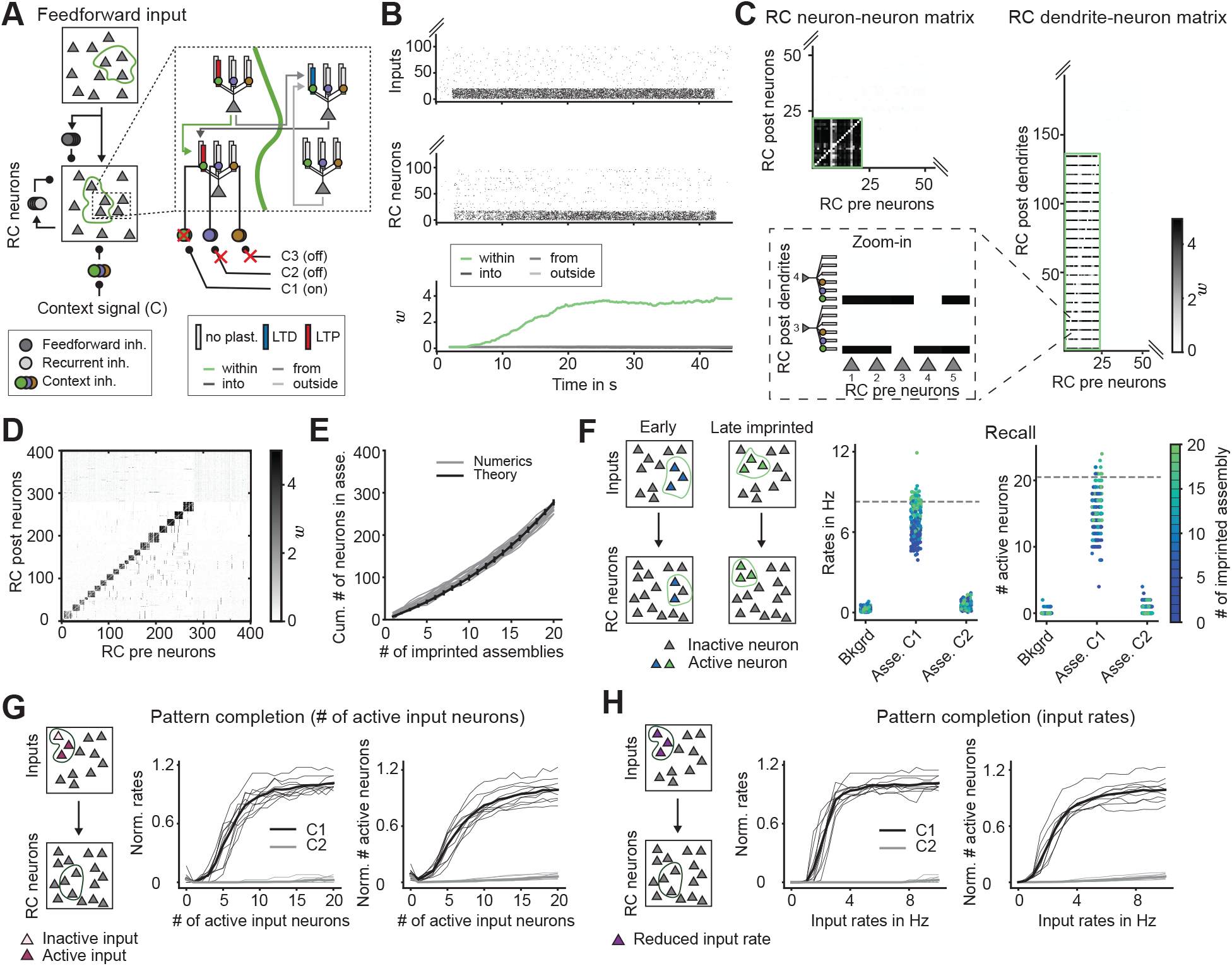
Nonlinear dendrites and inhibitory context lead to dendrite-specific assemblies. **A**. Schematic of feedforward inputs and recurrent (RC) network neurons. Inset: Schematic of connections and depolarization levels at different dendrites when context 1 (C1) is ‘on’. Two neurons are within (left) and two neurons are outside (right) of the assembly. **B**. Feedforward input spikes (top), recurrent neuron spikes (middle), and mean recurrent weights (*w*) over time. Weights are separated into groups corresponding to whether they are between neurons within the assembly (green line), from the assembly, into the assembly, and outside the assembly (gray lines). **C**. The representation of a single assembly through connectivity matrices. Sorted neuron-neuron (left) and neuron-dendrite (right) connectivity matrix. Grayscale indicates connection strength. Inset: Subset of presynaptic neurons and dendrites. **D**. Neuron-neuron connectivity matrix of the recurrent neurons after imprinting 20 assemblies one after another. Neurons are sorted in descending order, starting from the first imprinted assembly. **E**. The dependence of the cumulative total number of neurons that are part of any assembly on the number of imprinted assemblies can be predicted from the statistics of random connection probability (Methods). **F**. Recall in recurrent networks depends on the order in which assemblies were imprinted (left). Recall is measured via firing rates (middle) and number of active assembly neurons (right). Color indicates the order in which assemblies were imprinted (Asse. C1 - recall of an assembly in the learned context, Asse. C2 - recall of the same assembly in a different context, Bkgrd refers to the firing rate of non-assembly neurons). Recall is repeated 5 times for each assembly, and the dashed line represents the mean for the last imprinted assembly. **G**. Pattern completion in recurrent networks as a function of number of active input neurons measured via the firing rates of assembly neurons (middle) or number of active assembly neurons (right) in the context in which the assembly was learned (C1; black) or in a different context (C2; gray). The same 20 active neurons as the last imprinted assembly in panel F. **H**. Same as panel G as a function of the input firing rates. Input rate of 10 Hz corresponds to the recall in panel F.

Next, we investigated how our assembly framework can be used to form multiple assemblies by training the weights in the same context (C1). To determine how many assemblies can be stored, we sequentially activated non-overlapping groups of some number of (e.g. 20) feedforward inputs and projected them one-after-another (e.g. 20 groups in total) into the downstream recurrent area (Fig. 3D). Because of the random connectivity between the feedforward inputs and the recurrent neurons, it is unlikely that these non-overlapping feedforward inputs generate non-overlapping assemblies in the target area. Some neurons that have been part of a previously learned assembly might become recruited by a newly formed assembly, while other neurons in the recurrent area do not become part of an assembly because they never receive sufficiently strong excitatory inputs. For the choice of 20 groups with 20 input neurons each, ~ 3/4 of neurons in the recurrent downstream area became part of an assembly with later imprinted assemblies having more neurons (Fig. 3D). The number of assemblies and the assembly size can be well described from the statistics of random connection probability, in which earlier imprinted assemblies will have smaller assembly sizes (Methods, Fig. 3E).

A key feature of assembly learning is that those previously learned assemblies can be reliably reactivated—or ‘recalled’ (Hopfield, 1982; Vogels et al., 2011; Zenke et al., 2015). Therefore, we next quantified to what extent the re-activation of assembly neurons in the recurrent network by their respective inputs after all assemblies have been imprinted reflects the original assemblies imprinted by the same inputs during training. We found that reactivating assemblies learned early in the training process have weaker firing rates (~4-6 Hz) and fewer active assembly neurons (~10-18) compared to those learned late in the training process (~8 Hz firing rates and ~20 active assembly neurons) (Fig. 3F). The assemblies learned first are more affected by this reduction than the ones learned later, as their constituent neurons are likely to be recruited by newly learned assemblies. Nonetheless, both measures are clearly above the background activity of non-assembly neurons, suggesting that all learned assemblies can be reliably recalled. In addition, switching to a different context (C2) than the one used during training (C1) does not re-activate the assembly learned in the original context (C1) (Fig. 3F, compare Asse. C1 and Asse. C2). Therefore, recall of assemblies in our framework depends on the context in which those assemblies are learned, a feature that we later use to perform assembly computations.

Storing memories in strongly recurrently connected neuron groups, which represent the assemblies, is known to amplify weak stimuli (Peron et al., 2020) and enable partial stimuli to evoke a full assembly response – this has been termed pattern completion (Hopfield, 1982; Zenke et al., 2015; Vogels et al., 2011; Guzman et al., 2016). We examined pattern completion by testing the response in the recurrent network to changing the properties of the input neurons used during training. First, we quantified pattern completion by reducing the number of active inputs compared to those used during training (Fig. 3G). In the correct context (C1) for a given assembly, i.e., the context in which that assembly was learned, assembly firing rates and their number of active assembly neurons rapidly increase with the activation of more inputs (Fig. 3G, middle and right). Even reducing the number of active inputs by more than half compared to those used during training reliably activates the whole assembly in the recurrent network. This amplification is a direct result of the strong recurrent within-assembly weights in the recurrent network. When presenting a different context (C2), far fewer assembly neurons compared to context C1 are active and at much lower firing rates (Fig. 3G, gray lines). The network also exhibits pattern completion when reducing the input firing rate compared to those used during training (Fig. 3H).

In summary, we find that a large number of assemblies can be learned in the same context and reliably recalled, but only in the context in which they were learned. Due to the strong within-assembly connectivity, it is sufficient to weakly activate the input neurons or activate only a subset of them while still ensuring reliable assembly reactivation. Hence, the dendritic-specific assemblies learned by gating inhibition in the appropriate context can perform pattern completion and recall.

### Learning assemblies in multi-area networks without catastrophic forgetting

A prominent hypothesis puts forward assemblies as the basic computational units in the brain (Buzsáki, 2010; Byrne and Huyck, 2010; Herpich and Tetzlaff, 2019). Thus, we next investigated how the formed assemblies in our recurrent networks can be combined to perform computations. Specifically, we focused on stably learning projections from one area into another as well as learning associations across areas, as two types of ‘assembly calculus’ operations with a clear biological correlate underlying many neural computations (Papadimitriou et al., 2020; Papadimitriou and Friederici, 2022). For example, projections of assemblies are suggested to generate directed sequences in hippocampus (Holtmaat and Caroni, 2016), and assembly associations are useful to combine information from multiple assemblies into a new one (Pokorny et al., 2020). We focused on hierarchically organized multi-area networks without reciprocal feedback connections between them because many brain circuits are organized in a hierarchical manner (Harris et al., 2019; Siegle et al., 2021) and have inspired deep neural networks in artificial intelligence research (Yamins et al., 2014).

In our proposed architecture, the context in which an assembly is learned plays a crucial role for the calculus operations, as it defines which dendrites are gated ‘on’. To learn a projection in a given context C1 from one area (X) to a downstream area (Y), we activated a subset of neurons in area X which randomly project to a subset of dendrites in area Y. Our activity-dependent plasticity rule leads to the formation of a projection assembly with strong recurrent connections between those neurons with ‘on’-gated dendrites in area Y (Fig. 4A, B; light green).

**Figure 4.**
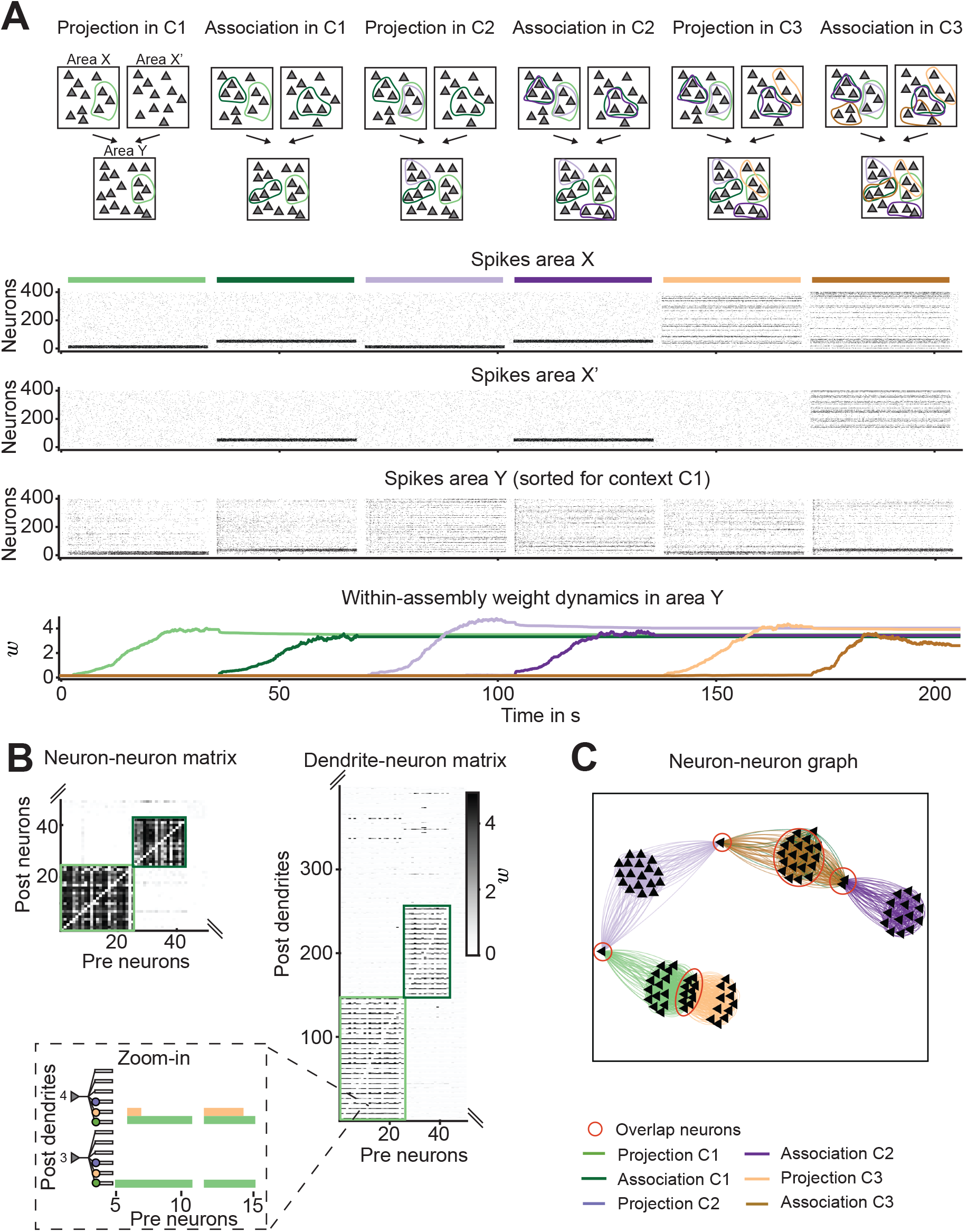
Learning without catastrophic forgetting. **A**. Top to bottom: Schematic of a network with three areas, spikes in area X, X’, and Y (spikes in area Y sorted to context C1), and mean within-assembly weight (*w*) change in different contexts (C1, C2, and C3). The firing rate of input spikes is 10 Hz. **B**. Neuron-neuron (left) and dendrite-neuron (right) connectivity matrix of area Y sorted according to context 1. Grayscale indicates connection strength. Inset: Subset of presynaptic neurons and dendrites with weights color-coded according to context. **C**. Neuron-neuron connectivity graph of area Y (only a subset of assembly neurons shown). Red circles indicate overlap neurons that are part of multiple assemblies.

In a second step, we aimed to learn an association of neurons from two different areas (X and X’) by forming an assembly in a common downstream area (Y) in the same context (C1) as the initial projection. Hence, we simultaneously coactivated a subset of neurons in area X and X’ (Fig. 4A, B; dark green). As before, synaptic plasticity led to the formation of an association assembly in area Y by potentiating the recurrent weights between those neurons with ‘on’-gated dendrites in area Y. Due to the presence of feedforward inhibition, we ensured that plasticity during both, projection and association, generated assemblies of similar size (Methods, Suppl. Fig. S2F, G).

A common problem with learning assemblies in recurrent networks is catastrophic forgetting, as previously learned assemblies become erased by new ones (Litwin-Kumar and Doiron, 2014; Zenke et al., 2015). Our proposed framework with dendrite-specific inhibitory gating provides a solution to this problem. We again took the same neurons in area X involved in the initial projection in context C1, but activated them in another context (C2). This activated a different subset of neurons in area Y due to the sparse and random feedforward connectivity and the distinct sets of ‘on’-gated dendrites for contexts C1 and C2 (Fig. 4A; compare light green and light purple). Therefore, synaptic plasticity formed a different projection assembly in area Y in context C2 by potentiating the recurrent weights between a new subset of neurons (Fig. 4C; compare light green and light purple). Following the same approach, a new association can also be learned in the new context (C2) (Fig. 4A, C; dark purple).

Finally, to test if our framework supports learning with overlapping assemblies, we deliberately selected neurons in area X’ that project to those dendrites in area Y, which are gated ‘on’ in a new context (C3) and belong to the assembly neurons of the initial projection in C1 (light green). Activating these neurons (Fig. 4A; light brown) leads to the formation of an assembly consisting of a large number of the same neurons as the assembly learned in the initial projection, despite a completely different set of presynaptic neurons in X (light green) and X’ (light brown). The same approach works for an association in context C3 (Fig. 4A; dark brown). Therefore, we find that the same neurons can be part of multiple assemblies through their multiple dendrites, even with a large overlap between the assemblies (Fig. 4B, C; light and dark brown). Crucially, the already learned assemblies in area Y are not forgotten, since previously learned synaptic weights are protected when learning new assemblies by changing context (Fig. 4A; bottom).

In summary, dendrite-specific gating of synaptic plasticity via context-dependent inhibition enables the learning of multiple overlapping assemblies – projections and associations – without forgetting. While these assemblies are defined in the typical sense with strongly mutually connected neurons (Miehl et al., 2023), the dendrite-specific gating by context-dependent inhibition ensures that each neuron participates in an assembly only with its ‘on’-gated dendrite in a given context. Hence, imprinting multiple of these non-overlapping assemblies can support ‘assembly calculus’ operations underlying many neural computations (Papadimitriou et al., 2020; Papadimitriou and Friederici, 2022).

### Assembly computations in a visual-auditory association task

We designed a visual-auditory association task to demonstrate the applicability of the ‘assembly calculus’ framework in a real-world example. The goal of the task is to distinguish four different stimuli, two letters and two numbers, by their neural representation in a downstream area. We chose the numbers “0” and “1” and the letters “O” and “l” because both pairs have very similar visual appearances (“0” and “O” vs. “1” and “l”), but differ in their phonetics ([Zee-Ro] and [Ou] vs. [Wun] and [El]). Hence, it is much more difficult to distinguish the stimuli by only relying on the visual modality. We represented the sensory stimuli in the form of assemblies, first in a visual and an auditory sensory area, followed each by a downstream (auditory and visual) area (Fig. 5A, left). We tasked the network to separate the two stimuli in a ‘concept’ area that is located downstream from the auditory and visual areas. We used two distinct contexts in the visual, auditory and concept areas to simulate the knowledge of perceiving letters vs. numbers.

**Figure 5.**
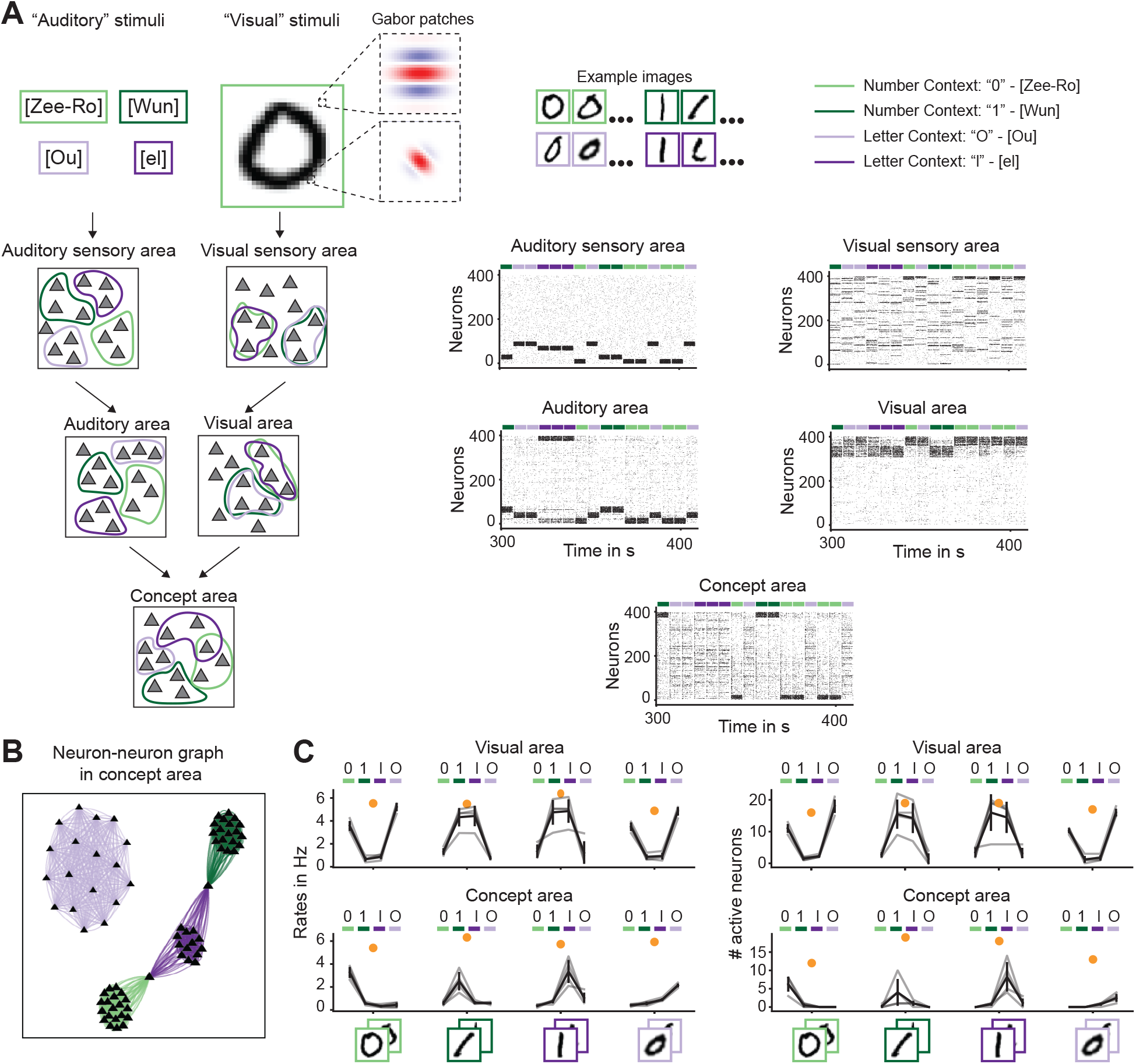
Assembly computations in a visual-auditory association task. **A**. Left: Schematic of visual-auditory association task, with two contexts: numbers (green) or letters (purple). Different auditory stimuli activate different assemblies in the auditory sensory area. Visual stimuli (taken from the extended MNIST dataset filtered by Gabor patches, see Methods) are more similar and activate similar assemblies in the visual sensory area. Right: Spiking activity in each area. Spikes are sorted according to the number context (green) in the auditory, visual, and concept areas. **B**. Neuron-neuron connectivity graph of the concept area. **C**. Quantification of pattern completion via assembly firing rates (left) or number of active assembly neurons (right) in the visual area (top) or the concept area (bottom) for each type of visual stimulus (“0”, “O”, “1”, and “l”) alone. Orange dots correspond to values when both auditory and visual stimuli are presented together. Grey lines are single images, black line is the mean across images.

Since all stimuli sound phonetically different, in the auditory sensory area, we assumed that each stimulus activates a potentially different assembly. In contrast, in the visual sensory area, similar stimuli (“0” and “O” vs. “1” and “l”) activate largely overlapping assemblies. To train the network with visual stimuli, we choose images from the extended MNIST dataset which were filtered by Gabor patches (Methods). We implemented a learning protocol using our proposed plasticity framework with dendrite-specific inhibition encoded by context (numbers vs. letters): we projected each stimulus from the two sensory areas into assemblies in the downstream areas, associating the corresponding visual and auditory representations of the same stimulus, as can be seen in the activity (Fig. 5A; right). Although largely overlapping assemblies in the visual area represent similar visual stimuli so they cannot be easily distinguished, due to the association with assemblies from the auditory area and in the presence of different contexts for letters vs. numbers, we found that they can be well separated in the downstream concept area, as reflected in non-overlapping assemblies (Fig. 5B).

After learning, we tested the ability of the network to separate potentially confounding visual stimuli in the downstream concept area. We presented a new visual image not used during training in the absence of the corresponding auditory stimulus in each of the two contexts, and measured the assembly firing rates and the number of active assembly neurons in the downstream auditory/visual areas and the concept area (Fig. 5C). Unsurprisingly, the downstream visual area fails to distinguish the two pairs of visually similar stimuli (“0” and “O” vs. “1” and “l”) (Fig. 5C, top). However, due to the formed associations, the concept area can separate the two visually similar stimuli even without the presentation of an auditory stimuli, as represented by the high firing rate and higher number of active neurons of the corresponding assembly (Fig. 5C, bottom). Even when only providing half of the input (e.g./ visual stimulus only) to the concept area, strong recurrent connectivity among assembly neurons amplifies the firing rates and activates the entire assembly - consistent with pattern completion.

In summary, learning overlapping assemblies without forgetting enables context-specific assembly computations in the form of projections and associations, which can be used to correctly separate ambiguous stimuli in a given context by relying on their dendritic-specific representations.

### Recall and pattern completion are enhanced in downstream areas

The recall due to strong recurrent connectivity can prove useful in preventing information from being degraded as it propagates across multiple hierarchically organized areas. To investigate if this mechanism operates in our networks, we imprinted assemblies in several consecutive multi-area projections (from area X to area Y to area Z) through context-gated plasticity (Fig. 6A). Given that neurons can drop out of an assembly without affecting the firing rates of the assembly (Fig. 3G), we wondered if this dropout might impair the recall of assemblies in multi-area projections. Hence, we silenced a fraction of the neurons of the learned assembly in area Y and tested recall in area Z while reactivating the original set of neurons in area X (Fig. 6B). We observed that despite decreased activity in area Y, assembly neurons in area Z remain highly active due to the strong recurrent within-assembly connections. Just like recall, pattern completion in area Z can be achieved when activating a subset of neurons in area X even if a fraction of neurons in area Y are silenced (Fig. 6C). Therefore, the strong recurrent within-assembly weights effectively implement a chain of activity amplification as information propagates across multiple hierarchical areas due to pattern completion. Our results demonstrate that multi-area projections of neural assemblies can lead to reliable recall in downstream areas despite disruptions in earlier, upstream areas.

**Figure 6.**
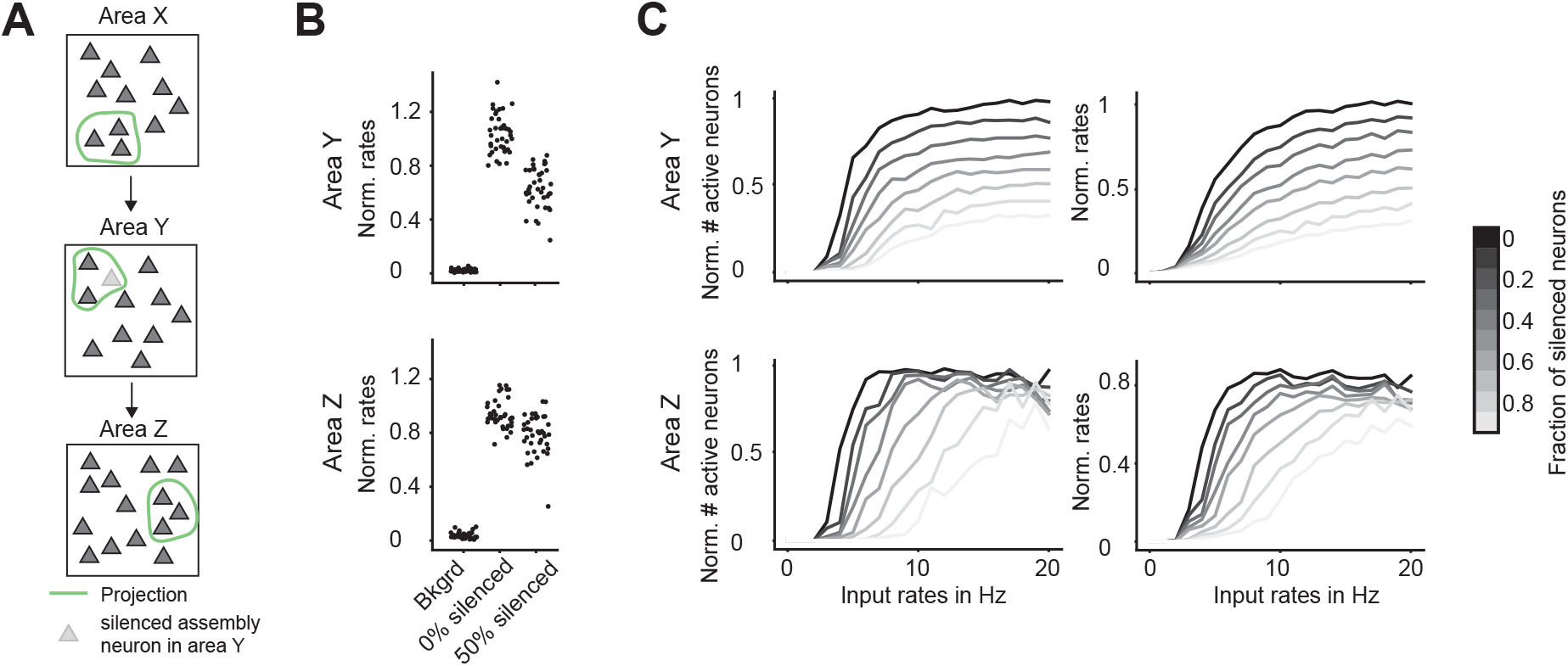
Enhanced recall and pattern completion in downstream areas despite disrupted recall in previous areas. **A**. Schematic of a network with three hierarchically connected areas X, Y, and Z. Assemblies are denoted by the green circled neurons which project across the areas. Shaded triangles denote silenced neurons in area Y. **B**. Quantification of recall for 0%, and 50% silenced assembly neurons in area Y (top) and area Z (bottom) via normalized assembly firing rates. ‘Bkgrd’ indicates background firing rates. **C**. Left: Quantifying pattern completion while silencing assembly neurons in area Y (gray-scale) via the normalized number of active neurons. Right: Same as the left panel for the assembly’s normalized firing rates in area Y (top) and area Z (bottom).

### Associative learning with existing assemblies

So far, we showed that our framework supports the learning of associations between different stimuli. However, these associations can be learned in different ways: ‘simultaneously’, where the assemblies are learned by activating them at the same time, or ‘sequentially’, where an association is learned utilizing existing assemblies. We compared how the identity of assembly neurons and pattern completion are implemented in these two types of associative learning as well as the case of learning two separate projections. Specifically, we studies three cases: (I) sequential projections of distinct assemblies, first from area X to Y to Z, followed by a second projection from area X’ to Y to Z; (II) simultaneous association of assemblies from areas X and X’ to Y to Z, and (III) sequential projection and association where the association of assemblies from areas X and X’ to Y to Z is learned on top of an existing projection from X to Y to Z (Fig. 7A). We quantified how learned assemblies in areas Y and Z differ by comparing cases I and II (Fig. 7B) and cases II and III (Fig. 7C) based on three measures: the overlap in assembly neuron identity (Venn Diagram), the distribution of input synapses onto assembly neurons (Synapse distribution), and the response of assemblies to partial activation of input neurons (Pattern completion).

**Figure 7.**
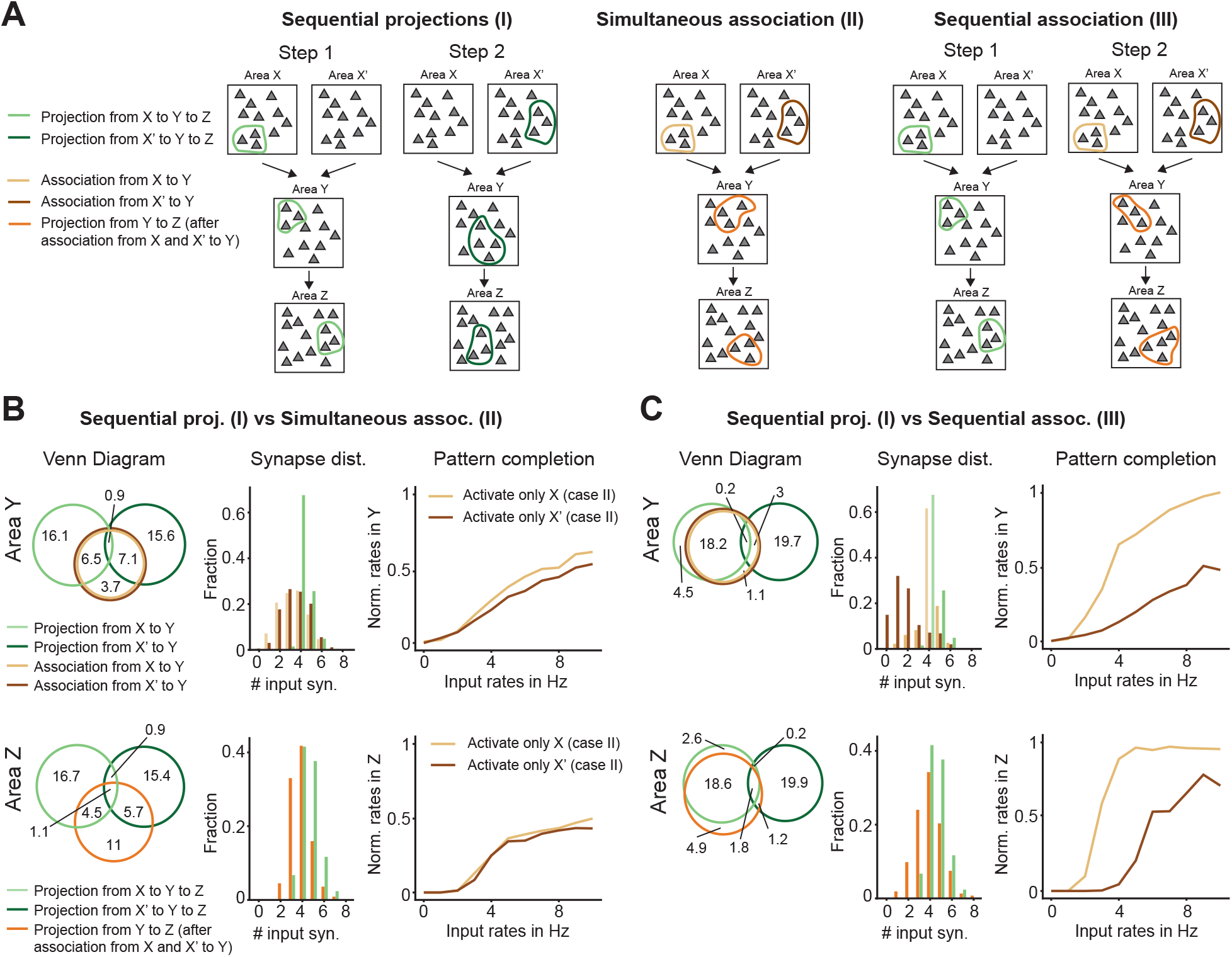
Associative learning with existing assemblies. **A**. Schematics of three different assembly formation scenarios across multiple areas: Sequential projection (I), simultaneous association (II), and sequential association (III). For the comparison across all cases, we used the same connectivity matrices across and within areas, activated the same inputs in areas X and X’, and used the same context. **B**. Compare sequential projection (I) versus simultaneous association (II). Left: Venn diagram of the average number of neurons in area Y (top) or area Z (bottom), which is either unique to each assembly (light/dark green and light/dark orange) or part of two or more assemblies (overlaps in the diagram). Middle: Input synapse distribution from area X or X’ to area Y (top) and from area Y to area Z (bottom). The color of the bars indicates the presynaptic assembly (X or X’) and the projection/association into the downstream assembly (Y or Z). (e.g. light orange in area Y shows the synapse distribution from the assembly in area X to the assembly in area Y in the case of simultaneous association, and the light green shows the case of the sequential projection). Right: Quantification of pattern completion via the normalized rates when activating only X (light orange) or only X’ (dark orange) for area Y (top) and area Z (bottom) for simultaneous association (case II). Rates were normalized to the response in area Y (top) and area Z (bottom) when activating both assemblies in X and X’ at the same time. **C**. Same as panel B, comparing sequential projection (I) versus sequential association (III).

First, we compared the assemblies formed in areas Y and Z via sequential projection (case I) or via simultaneous association (case II). A large subset of neurons in the simultaneous association assembly in area Y is the same as the neurons learned via the respective projection only (on average 6.5 for the light green and 7.1 for the dark green assembly), while a smaller subset of neurons (3.7) is unique to the assembly learned via the simultaneous association (Fig. 7B, top left). In the downstream area Z, the overlap decreases, with a smaller subset of neurons representing the learned association compared to learning projections alone (on average 4.5 for the light green and 5.7 for the dark green assembly), while the number of unique assembly neurons increases (11) (Fig. 7B, bottom left). Therefore, the assemblies learned via simultaneous association (case II) become more independent from assemblies learned via projections (case I) in more downstream areas.

We next quantified how input synapses onto assemblies in areas Y and Z differ between cases I and II. To form an assembly in area Y, neurons need to receive a sufficiently high number of active inputs to drive LTP. As previously shown (Fig. 2B), for the case of a projection from X to Y, most assembly neurons in the downstream area have to receive ≥ 4 highly active inputs from the upstream area (Fig. 7B, middle, light green bars in area Y and Z). For the case of simultaneous association, there are twice as many highly active assembly neurons in area Y since both X and X’ are active, in contrast to the case of a projection when only X is active. Therefore, the distribution of input synapses from area X and X’ onto the assembly in area Y in the case of simultaneous association (II) is broader compared to the distribution in the case of a sequential projection (I) (Fig. 7B, top middle, light and dark orange compared to light green). In the downstream area Z, the synapse distributions of the two cases become more similar again because the assembly in area Y (projection in case I and simultaneous association in case II) forms a projection to area Z in the two cases (Fig. 7B, bottom middle, orange compared to light green). When activating either only the inputs in area X or area X’ in the case of simultaneous association, the downstream assemblies in areas Y and Z fire reliably at ≈ 50% of the rates when activating X and X’ simultaneously, significantly above background firing rate (Fig. 7B, right, at 10 Hz input rates). Pattern completion is robust in both areas as observed before in Fig. 6.

Next, we compared the assemblies formed in areas Y and Z via sequential projection (case I) vs. sequential association (case III). In contrast to the comparison between I and II, most assembly neurons involved in cases I and III are similar, in particular, the neurons in areas Y and Z after learning the association are almost the same as the neurons learned only from a projection in X (on average 18.2 neurons in area Y and 18.6 neurons in area Z, Fig. 7C, top and bottom left). Therefore, when learning new associations with existing assemblies, most of the neurons that become part of the new assembly stay the same as the existing assembly. This is different from case II where the identity of assembly neurons differs, especially in downstream areas. The distribution of input synapses shows that most of the strong synapses onto the sequential association assembly in area Y come from previously imprinted area X (Fig. 7C, top and bottom middle). This leads to a more reliable pattern completion (higher firing rates) when activating only area X compared to activating only area X’ (light orange is higher than dark orange, Fig. 7C, top right). Moreover, when activating only X’, pattern completion is improved in the downstream area Z compared to area Y (dark orange is higher in the bottom than the top).

In summary, our framework allows us to learn two types of associations – either via simultaneously learned associations or by learning an association on top of existing projections. This suggests general principles of learning in the brain that might benefit from previously learned structures. While the learned assemblies differ in terms of which neurons and which synapses participate in intermediate areas along the hierarchy, their differences vanish in later downstream areas, where computational aspects like pattern completion become more reliable.

## Discussion

While assemblies have often been suggested as the basis for computations in the brain (Buzsáki, 2010; Eichenbaum, 2018; Huyck and Passmore, 2013; Yuste, 2015), few theoretical studies have shown how assemblies can be flexibly learned and combined to perform real-world complex computations. This has been mainly due to the challenge of learning overlapping assemblies without merging and the problem of catastrophic forgetting. Here, we propose two biologically-inspired mechanisms as a solution, specifically nonlinear dendritic compartments as the loci for learning, and inhibitory context-dependent gating allowing for dendrite-specific gating of synaptic plasticity via disinhibition (Fig. 2). Together with inhibitory control of assembly size, these mechanisms support the learning of stable but flexible assemblies (Fig. 3). These assemblies can be combined via projections or associations across multiple hierarchically organized areas without forgetting previously learned structures, and with assembly overlaps where the same neuron can participate in multiple assemblies through its different dendrites (Fig. 4). We exemplify assembly computations in a visual-auditory association task, in which we show how assembly computations can be used to learn associations of related stimuli across different sensory pathways and how they can be separated according to their representations in downstream areas (Fig. 5). We further show that assemblies can be reliably recalled, especially in downstream areas (Fig. 6) and that associative learning in these downstream areas may benefit from existing representations (Fig. 7).

The key underlying mechanism behind stable assembly formation and context-dependent gating in our framework is inhibition. We included inhibitory neurons with different roles in our modeling framework, in agreement with functions suggested by experimental studies. Recurrent inhibition prevents an assembly being learned from growing too large and taking over the entire network by suppressing excessive network excitation (Suppl. Fig. S2D, E). This mechanism is reminiscent of the ‘k-winner take-all’ concept (Kwon and Zervakis, 1995), where inhibition is proposed to keep roughly the same number of neurons active (Papadimitriou et al., 2020). Experimental support for this type of inhibition comes from the mouse hippocampus, where somatostatin-expressing (SST) inhibitory interneurons have been shown to control the size of co-active groups of neurons referred to as ensembles (Stefanelli et al., 2016). A second source of inhibition that acts in a feedforward manner in our model keeps the size of the assembly fixed when the excitatory input to the network changes (Suppl. Fig. S2F, G). This feedforward inhibition is consistent with that provided by parvalbumin-expressing (PV) interneurons (Tremblay et al., 2016). Changing the existing connectivity structure in our model implements learning, for which synaptic plasticity needs to be gated by decreased inhibition onto the dendrites, which are the loci of learning. Learning marks a departure of the model network from its baseline state where all dendrites are inhibited and synaptic plasticity is blocked. Learning occurs by selectively disinhibiting specific dendrites in a particular context. This disinhibitory mechanism is inspired by the disinhibitory influence of vasointestinal-peptide-expressing (VIP) interneurons onto SST interneurons, which together form the disinhibitory VIP-SST circuit. Hence, VIP interneuron input to SSTs provides a substrate for the context signal necessary for learning (Krabbe et al., 2019; Canto-Bustos et al., 2022). This disinhibition has been shown to enhance dendritic spikes (Gentet et al., 2012; Lovett-Barron, 2021) and flexibly route information (Yang et al., 2016). VIP interneurons are known to receive top-down inputs depending on the behavioral context of the animal and regulate population activity (Pi et al., 2013; Fu et al., 2014; Garcia Del Molino et al., 2017; Dipoppa et al., 2018).

Several prior computational studies have focused on how assembly structures can be learned with different plasticity rules (Clopath et al., 2010; Litwin-Kumar and Doiron, 2014; Zenke et al., 2015; Schulz et al., 2021; Miehl et al., 2023). However, many have faced the problem that any initial overlap between assemblies in terms of participating neurons usually leads to the merging or separation of the assemblies. A few notable example studies have demonstrated that assembly overlap is possible for specific choices of spike timing-dependent plasticity (Manz and Memmesheimer, 2023; Yang and Doiron, 2025), short-term plasticity (Fauth and van Rossum, 2019), or other mechanisms (Podlaski et al., 2025; Bergoin et al., 2025). A second related problem is forgetting previously learned assemblies where synaptic connections reinforcing one assembly decay while others grow as new assemblies are being learned. In our work, besides enabling learning at ‘on’-gated dendrites, the context signal also prevents synapses to ‘off’-gated dendrites from undergoing synaptic plasticity. This stabilizes the weights of already learned assemblies in different contexts, even if neurons of a previously learned assembly join a newly learned assembly (Fig. 3–5). Hence, the mechanism of context-dependent inhibition allows assemblies to overlap without merging or forgetting by sharing participating neurons through their different dendrites. This is similar to the idea that inhibition keeps memories in a quiescent state unless a context-dependent disinhibitory mechanism releases those previously learned memories (Barron et al., 2016, Barron et al., 2017). The proposal that inhibition might mediate a context signal has been implemented in several computational studies, studying the influence of surround information on visual computation (Voina et al., 2022), increasing capacity in memory networks (Podlaski et al., 2025), multitask and transfer learning (Wybo et al., 2023), and showing that sparse inhibitory context projects the network activity onto unique neural subspaces (Lehr et al., 2023).

Furthermore, our approach is similar to a recent proposal that error signals between sensory and expected information during learning are encoded in the dendritic excitatory/inhibitory balance. In these studies, top-down feedback signals arriving at the dendrites can disrupt this balance, gate local plasticity and, therefore, lead to the re-learning or updating of synaptic weights according to the error signal (Sacramento et al., 2018; Rossbroich and Zenke, 2025; Galloni et al., 2025). The contextual signal that gates dendrite-specific plasticity in our study could also be interpreted as such an error signal.

In artificial neural networks (ANNs), catastrophic forgetting has been known for a long time to hinder sequential learning (McCloskey and Cohen, 1989). Many different solutions have been suggested to counteract forgetting in ANNs, like metaplasticity (Jedlicka et al., 2022), gating via inhibition (Masse et al., 2018; Sezener et al., 2021; Tilley et al., 2023), dendritic compartments (Chavlis and Poirazi, 2021; Pagkalos et al., 2024), elastic weight consolidation (Kirkpatrick et al., 2017), neuromodulatory systems (Mei et al., 2022, 2025), and many more (Parisi et al., 2019; Zenke and Laborieux, 2024). Our model proposes the biologically motivated mechanisms of dendrite-specific gating via disinhibitory context pathways to counteract forgetting, which might be one of multiple strategies to achieve lifelong learning (Hadsell et al., 2020; Kudithipudi et al., 2022). Interestingly, the idea bears resemblance to a recent approach, where an additional module protects already learned connections in an artificial neural network by ‘rotating’ the inputs (Zeng et al., 2019).

In conclusion, this study presents a novel approach to assembly computations with two key biologically inspired mechanisms that put forward dendrites as the main place of learning: nonlinear dendritic compartments and inhibitory context-dependent gating. Our model demonstrates how assemblies can be flexibly learned and combined without merging or catastrophic forgetting. The important new assumptions in this framework align with multiple experimental findings on different roles of inhibition, offering many insights into complex brain computations that underlie flexible and stable learning. Our work also opens new avenues for how to make artificial neural networks that can do useful computations more biologically plausible and hence bridges the gap between biological realism and computational functionality in neural circuit modeling.

## Methods

### The excitatory neuron model

The neuron model is a multi-compartment model adapted from Yang et al. (2016) and features *D* = 6 dendritic compartments that are each coupled to the soma.

#### The soma

The soma is a leaky integrate-and-fire compartment whose voltage dynamics follow:

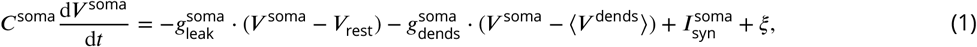

where *V* ^soma^ is the potential of the somatic compartment, *C*^soma^ = 50 pF is the membrane capacitance, 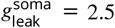 is the leak conductance and the conductance 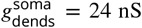 determines the coupling strength between all dendritic and the somatic compartment. *V*_rest_ = −70 mV is the resting potential and ⟨*V*^dends^⟩ describes the average potential of all 6 dendrites. In addition, the soma receives inhibitory gamma-aminobutyric acid (GABA) input via 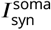 and somatic noise *ξ* is modeled as a Gaussian random variable with mean and standard deviation 110 ± 7 pA to produce a baseline spiking behavior and model excitatory and inhibitory background inputs. Whenever the somatic potential *V* ^soma^ exceeds the threshold *V*_thresh_ = −50 mV, the soma elicits a spike, and the somatic potential is set to *V*_reset_ = −55 mV. During refractory time *τ*_ref_ = 2 ms following the spike, *V* ^soma^ can change, but crossing the threshold again does not evoke a spike.

The soma features inhibitory synapses modeling the linear dynamics of GABA_A_ channels:

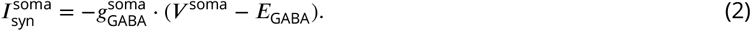

The reversal potential for GABAergic synapses *E*_GABA_ is set at −75 mV. The synaptic conductance 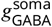 increases by 8 nS whenever a presynaptic spike arrives and decreases exponentially with time constant *τ*_GABA_ = 20 ms:

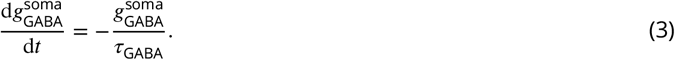

#### The dendrites

The dendrites are leaky integrator compartments, each with potential *V* ^dend^ following

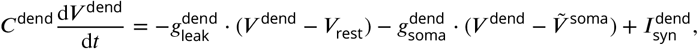

where *C*^dend^ = 20 pF is dendritic membrane capacitance, 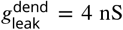 is the leak conductance and *V*_rest_ = −70 mV is the resting potential. The conductance 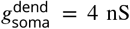 determines the coupling strength of the dendrite to the soma. 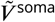 is an auxiliary somatic potential, whose dynamics are identical to *V* ^soma^ (Eq. 1 with exact same noise *ξ*) except that it does not spike. The dendrites feature inhibitory GABA receptor synapses and excitatory synapses consisting of *α*-amino-3-hydroxy-5-methyl-4-isoxazolepropionic acid (AMPA) and non-linear N-methyl-D-aspartate (NMDA) receptor channels:

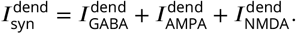

The currents 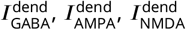 represent total currents flowing through the respective channels. The GABA and AMPA receptors follow linear dynamics analogous to (Eq. 2) and (Eq. 3):

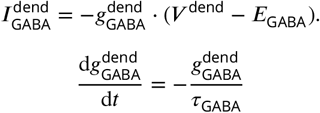

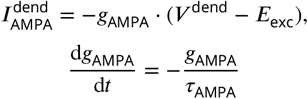

The reversal potential for excitatory synapses is *E*_exc_ = 0 mV and the time constant is *τ*_AMPA_ = 2 ms. With each presynaptic spike, the conductance *g*_AMPA_ is increased by 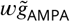, where 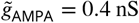 and the weight factor *w* may change due to synaptic plasticity. The conductance of GABAergic synapses *g*_GABA_ is increased with each presynaptic spike by a factor 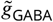, which depends on the identity of the inhibitory synapse (see below for the description of different types of inhibition). NMDA receptors introduce a non-linear excitation mechanism which includes a voltage-dependent magnesium block *f*_Mg_ and a saturating gating variable *s*_NMDA_:

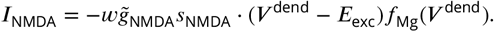

The conductance 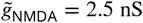. The weight *w* is shared with AMPA receptors within the same excitatory synapse. The magnesium block is modeled using a sigmoid function

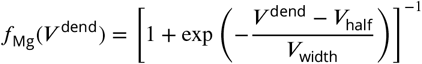

with its inflection point at *V*_half_ = −19.9 mV and slope governed by *V*_width_ = 12.48 mV. The NMDAR gating variable dynamics follow:

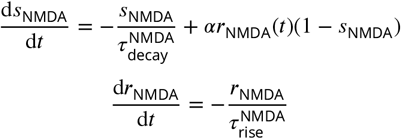

with 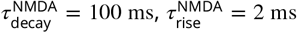, and *α* = 0.3 ms^−1^. Whenever a presynaptic spike occurs, *r*_NMDA_ is set to 1. In the case of linear NMDA receptors, *f*_Mg_(*V* ^dend^) = *V* ^dend^ (Supplementary Figure S1).

### Homosynaptic plasticity

We implemented a phenomenologically derived synaptic plasticity rule, the so-called voltage rule (Clopath et al., 2010; Bono and Clopath, 2017). The rule is a form of spike-timing-dependent plasticity that considers the timing of presynaptic spikes and the postsynaptic membrane potential (here the potential of a dendrite). The amount by which the synaptic weight *w* changes follows

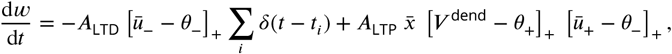

where 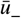 and 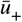 is the dendritic potential, delayed and low-pass-filtered with different time constants:

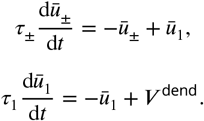

The []_+_ operator indicates rectification, that is [*λ*]_+_ = *λ* for *λ* > 0 and [*λ*]_+_ = 0 otherwise. The spike trace 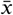 follows

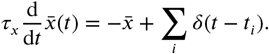

The Dirac *δ* represents presynaptic spikes occurring at times *t*_*i*_. When a spike arrives at a plastic synapse at time *t*_*i*_ and the dendrite has been sufficiently depolarized 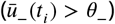 the synaptic weight gets immediately depressed by 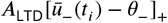. The synaptic weights may also undergo potentiation at all times when 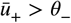 and *V* ^dend^ > *θ*_+_. The potentiation degree depends on the depolarization level and the time since the last presynaptic spikes. The amplitude parameters are *A*_LTD_ = 4 mV^−1^s^−1^ and *A*_LTP_ = 1.4 ⋅ 10^−3^ mV^−2^s^−1^, the threshold potentials are *θ*_−_ = −65 mV and *θ*_+_ = −30 mV, the time constants are *τ*_−_ = 15 ms, *τ*_+_ = 45 ms, *τ*_1_ = 5 ms and *τ*_*x*_ = 20 ms. We note that the values of *θ*_−_ and *θ*_+_ were slightly different from the ones used in (Bono and Clopath, 2017) to tune down synaptic depression. Both, feedforward and recurrent excitatory synapses undergo synaptic plasticity, and synaptic strength is bound within a specific range. For recurrent synapses the bounds are *w*_min,REC_ = 0 and *w*_max,REC_ = 5. For feedforward synapses, the maximum bound is given by *w*_max,FF_ = 26. The minimum bound is calculated for each dendrite individually to ensure that each dendrite can achieve a stable configuration when being part of an assembly:

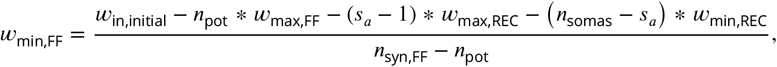

with the total initial synaptic weight of the incoming synapses *w*_in,initial_, the average number of potentiated feed-forward dendrites *n*_pot_, the number of somas in the area *n*_somas_, the average assembly size *s*_*a*_ and the number of feedforward connections *n*_syn,FF_.

### Heterosynaptic plasticity

Besides weight changes induced by the voltage rule, excitatory synapses on each dendrite were subject to a normalization procedure. Every 5 ms, each excitatory synapse (both recurrent and feedforward) on a dendrite *d* is potentiated or depressed by 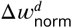 depending on the weights of other excitatory synapses *j* on the same dendrite.

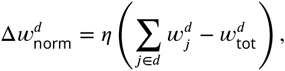

with *η* = 0.0025. The equilibrium value 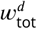 is the sum of initial weights of all feedforward and recurrent excitatory synapses incoming to dendrite *d*.

### Single neuron simulations

For the simulations exemplifying the working of a single neuron (Fig. 2), we built neurons with six dendrites. Each neuron is simulated with a combination of a set number of excitatory (active) synapses and a certain firing rate of the inhibitory neuron (as shown in Fig. 2C). Each neuron receives 399 weak (*w* = *w*_0_) excitatory connections representing recurrent input from other recurrent neurons and 32 stronger excitatory connections (*w* = *w*_ff_) representing the feedforward drive. The excitatory connections project synapses to all six dendrites of a neuron. The feedforward synapses receive random Poisson spikes at rate 0.1 Hz, except a set number of connections called active (Fig. 2C), which spike at rate 10 Hz. Additionally, each neuron’s dendrites receive GABAergic synapses from their inhibitory context neuron, which sends Poisson spikes at the indicated rates (Fig. 2C; constant across each neuron).

### Network simulations

For simulations that involve multiple networks (areas) of neurons, each area consists of 400 neurons. Each soma projects excitatory synapses to all dendrites of the network apart from its own with an initial weight of *w* = 0.1. Input to each area either comes from other areas (e.g. area Y gets input from areas X and X’ in Fig. 4) or two population 400excitatory neurons with Poisson neurons firing at the background rate *r*_bg_ = 0.1 Hz and a subset of 20 neurons with increased spiking rate of *r*_active_ = 10 Hz. The input neurons project to the network randomly with probability *p*_*FF*_ = 0.081. These connections underly homo- and heterosynaptic plasticity.

Apart from the recurrent and feedforward excitatory synapses, the neurons in the network receive inhibition from different sources, namely context inhibition, feedforward inhibition, and recurrent inhibition.

#### Context inhibition

Context is implemented as a population of inhibitory neurons, with each inhibitory population connecting to one dendrite of each neuron in the network. The context neurons inhibit the dendrite with a GABAergic synapse of static weight with the conductance increment 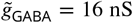 (see above). While active, a context neuron generates spikes according to a Poisson distribution with the mean rate *r*_context_ = 400 Hz. The disinhibition of dendrites by the context signal is implemented by setting the firing rates of the respective context neurons to 0 Hz.

#### Feedforward inhibition

Feedforward inhibition is implemented via a population of 400 inhibitory neurons projecting synapses to randomly selected dendrites in the network with a static weight 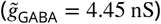. Each dendrite in the network receives a connection from one and only one feedforward inhibitory neuron. The feedforward inhibitory neurons are modeled as Poisson neurons with a rate depending on the activity of excitatory feedforward inputs to the network:

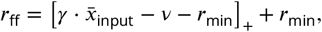

where *γ* = 0.34 Hz, *v* = 38.8 Hz, and 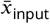 is a trace of activity of the excitatory inputs with a time constant *τ*_input_ = 750 ms:

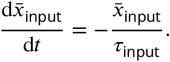

Whenever any feedforward excitatory neuron spikes, 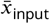 is increased by 1. The rectification []_+_ is shifted to keep the firing rate not less than the minimal value of *r*_min_ = 36 Hz. Feedforward inhibition leads to a fixed assembly size when changing the number of excitatory inputs (Suppl. Fig. S2F, G).

#### Recurrent inhibition

Recurrent inhibition controls excessive excitation in the network. For each excitatory neuron in the recurrent network, we computed a trace of its activity *x*_act_ low-pass-filtered with time constant *τ*_act_ = 750 ms:

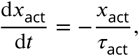

with *x*_act_ being increased by 1 whenever the neuron spikes. Each area receives input from 400 recurrent inhibitory neurons. These inhibitory neurons receive random connections from the somas in the network with probability *p*_rec1_ = 0.15. Whenever at least *n*_act_ = 8 of its presynaptic excitatory neurons become activated enough (their *x*_act_ > 2.2), the recurrent inhibitory neuron starts spiking in a Poisson manner at a rate *r*_rec_ = 60 Hz. The recurrent inhibitory neurons project randomly to the dendrites of the network with probability *p*_rec2_ = 0.06 and a static weight such that 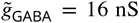. This setup allows the recurrent inhibition to control activity and plasticity based on the network activity, preventing excessive excitation (Suppl. Fig. S2D, E).

In addition, the somas of excitatory neurons receive GABAergic connections of static weight 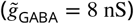 from a population of 400 inhibitory neurons. These neurons are connected one-to-one to the network somas, and they spike according to the Poisson distribution with the rate *r*_som_ = 40 Hz.

#### Sensory Stimulus Encoding

To model how different sensory areas process stimuli, we implemented distinct encoding strategies for the visual and auditory inputs (Fig. 5). In both cases, stimuli were represented by assemblies of 20 co-active neurons, but with different patterns of overlap between similar stimuli.

## Visual Stimulus Processing

We selected four characters from the Extended MNIST (EMNIST) dataset (Cohen et al., 2017): two pairs of visually similar characters (“0”/”O” and “1”/”l”). For each character, we randomly selected 15 different sample images. These grayscale images (28×28 pixels) were processed using Gabor filtering to simulate early visual processing.

We created a bank of 400 Gabor filters parameterized by:

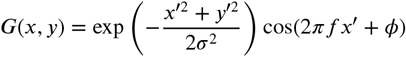

where (*x*′, *y*′) = (*x* cos *θ* + *y* sin *θ*, −*x* sin *θ* + *y* cos *θ*) represents coordinates rotated by angle *θ*. We systematically varied the Gabor parameters: orientation *θ* ∈ 0^°^, 45^°^, 90^°^, 135^°^, phase *ϕ* ∈ 0^°^, 180^°^, frequency *f* ∈ 1, 1.5, and scale *σ* ∈ 0.2, 0.4.

Each Gabor filter was assigned a random center position (*x, y*) in the grid, while ensuring equal coverage of the whole image. The response of each filter was computed by convolving the image with the filter. To generate a sparse input pattern for a given stimuli, we assigned all inputs a base value of 0.1 Hz to mimic the baseline firing rate. The ids of the weakest filter responses were set to 0 Hz, while strongest filter responses were scaled to keep their proportions but achieve an average value of 10 Hz (values are clipped if they exceed 14 Hz during this process). The main purpose of this filtering process was to preserve the visual similarity between character pairs, resulting in partially overlapping assembly activations for similar characters (e.g., “0” and “O”), while generating similar input patterns that were used during the rest of the paper.

## Auditory Stimulus Processing

For the auditory inputs, we implemented a direct mapping where each stimulus type was associated with a distinct subset of neurons. This mapping ensured that phonetically distinct stimuli activated non-overlapping sets of neurons in the auditory sensory area. Each auditory index corresponded to a specific group of 20 consecutive neurons in the feedforward population (e.g., neurons 0-19 for “1”, 20-39 for “l”, etc.). By implementing these contrasting encoding strategies, we captured a key feature of biological sensory processing: visual stimuli with similar appearances produce overlapping neural representations, while auditorily distinct stimuli (different phonemes) activate largely separate neural populations.

### Analysis

#### Identification of Neuronal Assemblies from Firing Rates and Synaptic Connectivity

To extract neurons that make up a neural assembly, we employed a two-stage clustering procedure that first uses firing rate statistics and subsequently refines the selection based on recurrent synaptic connectivity. First, we applied k-means clustering with k=2 to partition the neurons into two groups. The cluster whose centroid had the highest value was identified as the high firing rate group that forms the initial candidate assembly. In addition, we included the 10 neurons with the highest firing rate of the low firing rate group. Then, we applied k-means clustering with k=2 on the weight matrix that was reduced to contain only the candidate assembly neurons but included the summed input weights as an additional feature. Finally, we computed the mean intra-cluster connectivity for each cluster, and identified the cluster that exhibits the higher mean connectivity as the neuronal assembly.

#### Theoretical Calculation of Assembly Storage Capacity

In our model, the feedforward projections into an area are randomized with probability p=0.081. With 400 neurons and a target assembly size of 20, there are at max 20 non-overlapping inputs possible.

Statistically, it is unlikely that these non-overlapping inputs generate non-overlapping assemblies in the target area. We compared the assembly sizes of subsequently imprinted assemblies in a single context (Fig. 3E) with the theoretically determined sizes purely based on the statistics of how neurons in the target area are recruited.

We therefore repeated the following procedure 1000 times to sample the possible distributions of assembly sizes:

1. Initialize 20 empty assembly groups *A*_1_, *A*_2_, …, *A*_20_.
2. Sample an assembly size *s*_*i*_ from the empirical distribution of assembly sizes as observed in 500 network simulations with recurrent inhibition enabled (see Fig. S2E).
3. Randomly select *s*_*i*_ neurons from the pool of all neurons to form assembly *A*_*i*_.
4. Remove neurons from assemblies *A*_*j*_ with *j* < *i* for any neuron that appears in the new assembly *A*_*i*_, modeling the recruitment of neurons from earlier assemblies.
5. Calculate the cumulative number of neurons 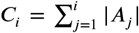 for each *i* from 1 to 20, where |*A*_*j*_| denotes the number of neurons in assembly *A*_*j*_ after all overlaps are accounted for.

This procedure captures the statistical properties of assembly formation in our network, particularly the effect of new assemblies recruiting neurons from previously formed assemblies. The calculated theoretical curve closely matches the results observed in the full network simulations (Fig. 3E), validating our understanding of the mechanisms that constrain assembly storage capacity.

## Code availability

All numerical computations were performed in Python using the Brian 2 simulator (Stimberg et al., 2019). The code will be available on https://github.com/comp-neural-circuits/contextual_gating after publication.

## Acknowledgements

We thank all members of the ‘Computation in Neural Circuits’ group and Dylan Festa, Filippo Kiessler, Satchal Postlewaite, and Joey Stawyskyj for useful discussions and comments on the manuscript. This project has received funding from the European Research Council (ERC StG NeuroDevo, Grant agreement No. 804824 to JG), the Human Frontier Science Program Organization (RGP0062/2021 to JG), and a Human Frontier Science Program Postdoctoral Fellowship (LT0005/2024-L to CM).

## Supplementary Text 1

### Necessity for additional mechanisms to stabilize learning in the recurrent circuit

To more robustly learn assembly structures, we assumed that the feedforward connections are also plastic. Synapses of highly active feedforward inputs increase their strength onto the ‘on’ gated dendrites in the recurrent circuit, while low active feedforward inputs decrease their strength (Suppl. Fig. S2A).

To control the size of assemblies and avoid enlarged assemblies, we added heterosynaptic normalization and two types of inhibition to our model. To prevent synaptic weights into the assembly from increasing due to synaptic plasticity, synapses are also subject to a dendrite-specific normalization mechanism, maintaining the total synaptic input strength on a given dendrite at a target value (Methods). This type of plasticity is known as heterosynaptic plasticity (Chistiakova et al., 2015; White et al., 1990; Royer and Paré, 2003; El-Boustani et al., 2018), as it affects synapses that are not directly activated by presynaptic inputs but are on dendritic branches where other synapses undergo synaptic plasticity. Therefore, when some input weights undergo LTP due to activity-dependent plasticity, others on the same dendrite undergo LTD due to heterosynaptic dendrite-specific normalization. This mechanism prevents the uncontrolled growth of newly formed assemblies (Suppl. Fig. S2B, C). Specifically, without heterosynaptic plasticity the weights from non-assembly neurons into the assembly grow without bound (Suppl. Fig. S2B, top), which eventually leads to enlarged assemblies where almost each neuron in the recurrent network projects onto the assembly neurons (Suppl. Fig. S2C). Also, heterosynaptic plasticity avoids all the connections between non-assembly neurons (outside connections) to decay (Suppl. Fig. S2B, bottom). This happens because many non-assembly neurons are in the LTD regime, hence those dendrites would lose all of their inputs without heterosynaptic normalization.

As assemblies form through the strengthening of recurrent connections between neurons, activity in the recurrent network increases, which substantially increases the number of dendrites with potentiated synapses, and therefore increases the size of the assembly over time (Suppl. Fig. S2D, left). This feedback excitation is a common problem in recurrently connected networks experiencing Hebbian plasticity (Litwin-Kumar and Doiron, 2014; Zenke et al., 2015; Miehl and Gjorgjieva, 2022). To control assembly size due to the positive feedback excitation in the recurrent network, we implemented recurrent inhibition (Suppl. Fig. S2D, right). Unlike the inhibition providing context to each dendrite by gating it ‘on’ or ‘off’ and determining its participation in an assembly, recurrent inhibition integrates local excitatory activity in the network and targets a random subset of dendrites (Methods). Indeed, assemblies with recurrent inhibition can be well confined and regulated (Suppl. Fig. S2E).

To regulate assembly size when changing the number of active feedforward inputs, we also implemented feed-forward inhibition (Methods). Without feedforward inhibition, increases in the number of feedforward inputs lead to increases in assembly size (Suppl. Fig. S2F, left). This is because by increasing the number of feedforward inputs, more dendrites will receive a sufficient number of inputs to push them into the LTP regime (Suppl. Fig. S2G, bottom). Adding feedforward inhibition, in which the amount of inhibition is proportional to the excitatory drive, keeps the assembly size fixed independent of the number of feedforward input neurons (Suppl. Fig. S2F, right). This follows because the number of highly active inputs needed to shift the dendrite into an LTP regime (i.e. the LTD/LTP threshold) is increased for more input neurons, keeping the number of dendrites in the LTP regime constant (Suppl. Fig. S2G).

## Supplementary Figures

**Figure S1.**
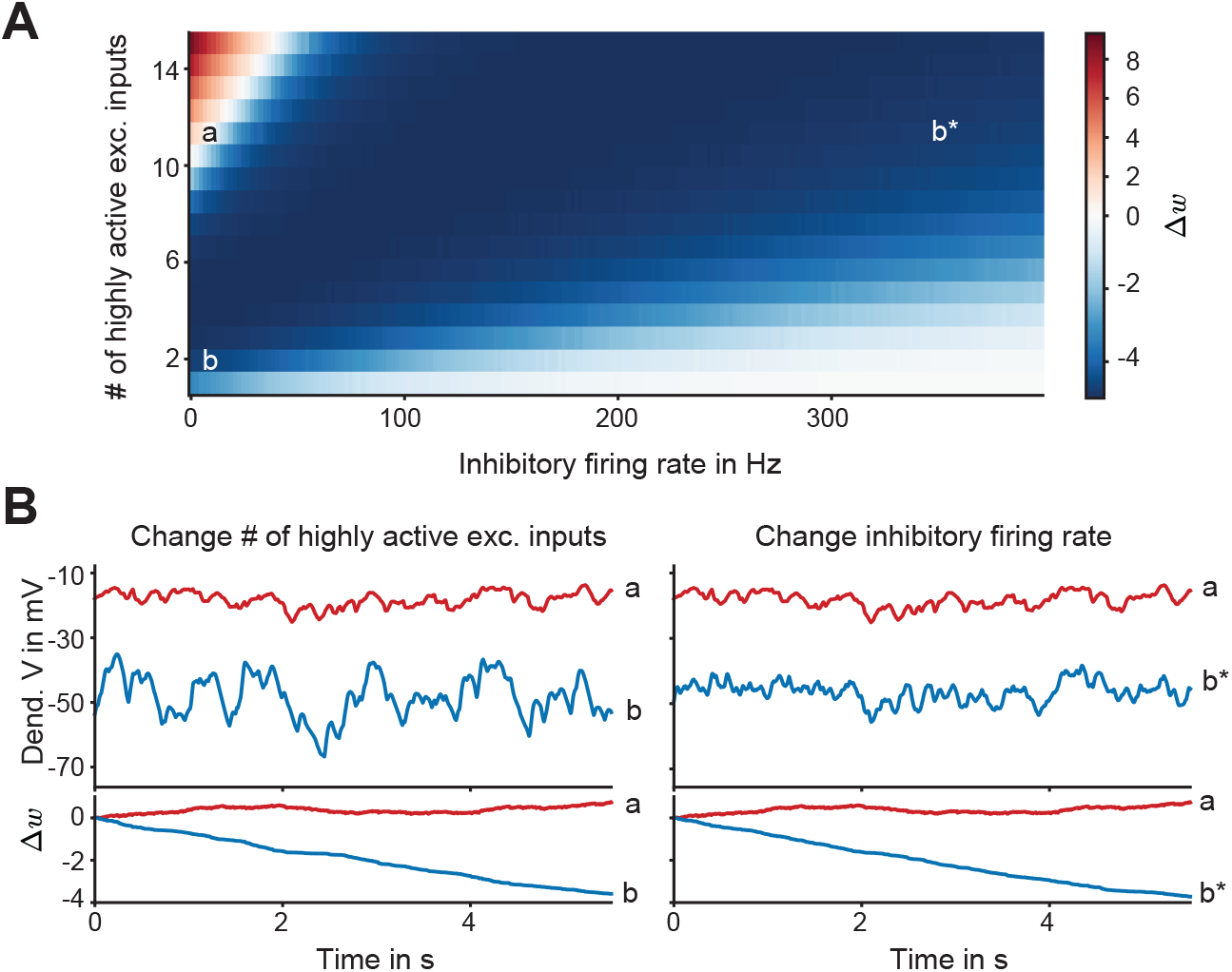
Weight changes in the case of linear NMDA receptors. **A**. Mean synaptic weight change Δ*w* as a function of the number of highly active excitatory synapses and the firing rate of inhibitory neurons. Long-term depression (LTD) is indicated by shades of blue, and long-term potentiation (LTP) by shades of red. **B**. Left: Filtered dendritic voltage *u*_+_ (top), and mean synaptic weight change Δ*w* (bottom) for a different number of highly active excitatory inputs (11 – red, 1 – blue). Letters (a,b) correspond to cases in panel A. Right: Same as left, for changing the inhibitory firing rate (10 Hz – red, 300 Hz – blue). Letters (a,b*) correspond to cases in panel A.

**Figure S2.**
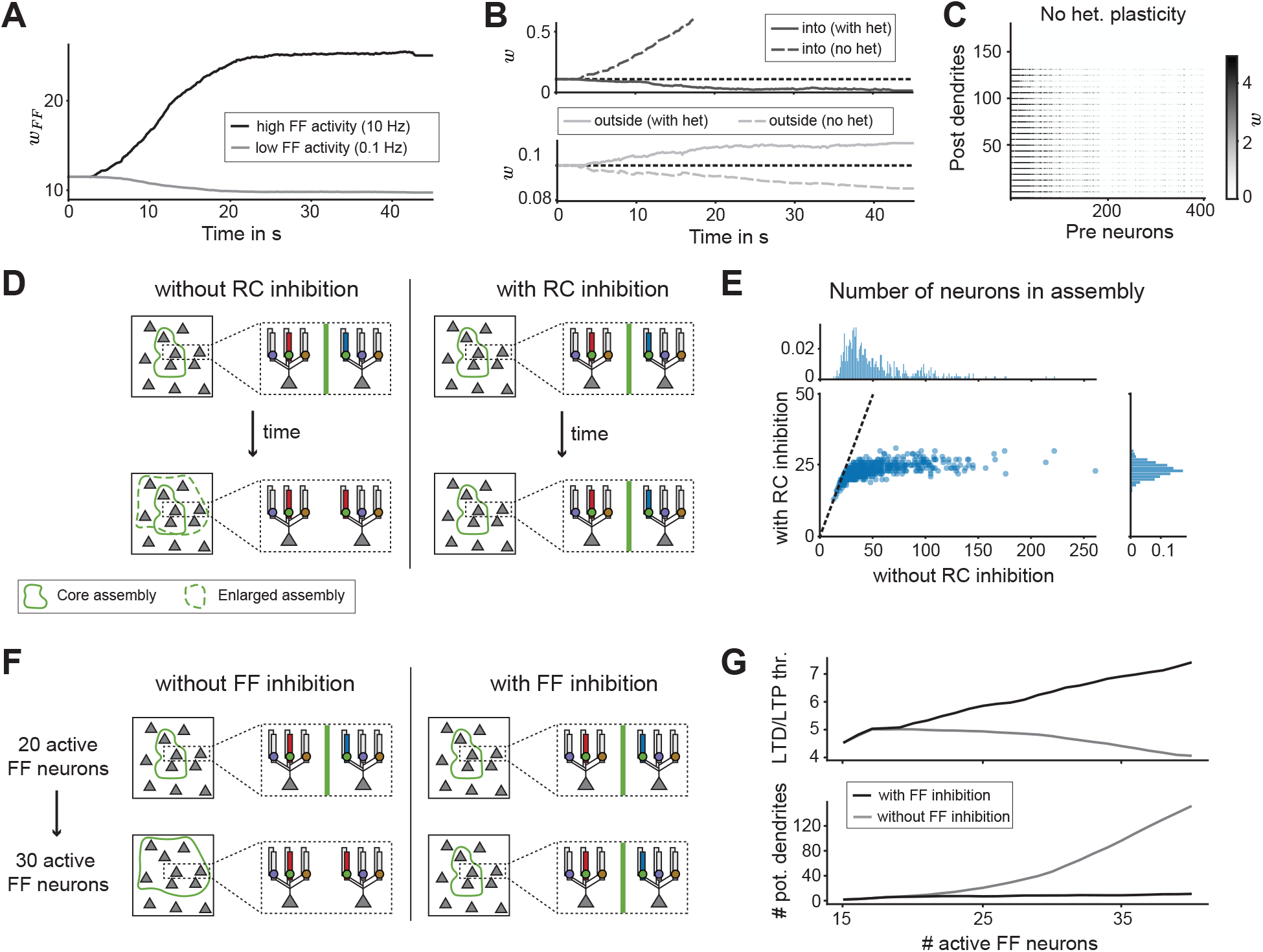
Necessity for additional mechanisms for stable learning in the recurrent circuit. **A**. Mean feedforward weight *w*_*FF*_. Weights are split into feedforward weights onto assembly neurons from highly active FF neurons (black line, 10 Hz) and low active FF neurons (gray line, 0.1 Hz). **B**. Mean recurrent weight *w* of into (top), and outside (bottom) assembly synapses without (dashed line) and with (solid line) heterosynaptic plasticity. The dotted black line represents the initial weight strength. **C**. Sorted neuron-dendrite connectivity matrix without heterosynaptic plasticity. **D**. Schematic of the effect of recurrent (RC) inhibition. Left: Without RC inhibition, newly formed assemblies continuously increase in size. Right: RC inhibition can control the size of the assembly in the recurrent network. **E**. Number of neurons in an assembly with and without recurrent inhibition. **F**. The effect of adaptive feedforward (FF) inhibition. Left: Without FF inhibition, increasing the number of active feedforward inputs recruits bigger assemblies. Right: With FF inhibition, increasing the number of active feedforward inputs leads to the formation of assemblies of the same size. **G**. The number of highly active inputs needed to shift the dendrite into an LTP regime (i.e. the LTD/LTP threshold, top) and number of dendrites in the LTP regime (bottom) as a function of the number of active presynaptic neurons with (black) and without (gray) feedforward inhibition.

